# Revealing the causative variant in Mendelian patient genomes without revealing patient genomes

**DOI:** 10.1101/103655

**Authors:** Karthik A. Jagadeesh, David J. Wu, Johannes A. Birgmeier, Dan Boneh, Gill Bejerano

**Affiliations:** Department of Computer Science, Stanford University; Department of Electrical Engineering, Stanford University; Department of Developmental Biology, Stanford University; Department of Pediatrics (Medical Genetics), Stanford University

## Abstract

Given the rapidly growing utility of critical health information revealed in the human genome, secure genomic computation is essential to moving forward, especially as genome sequencing becomes commonplace. We devise and implement proof-of-principle computational operations for precisely identifying causal variants in Mendelian patients using secure multiparty computation methods based on Yao’s protocol. We show multiple real scenarios (small patient cohorts, trio analysis, two hospital collaboration) where the causal variant is discovered jointly, while keeping up to 99.7% of all participants’ most sensitive genomic information private. All similar operations performed today to diagnose such cases are done openly, keeping 0% of participants’ genomic information private. Our work will help usher in an era where genomes can be both utilized and truly protected.

## Introduction

Rare diseases affect 1 in 33 babies. Exome and genome sequencing have revolutionized the diagnosis of thousands of rare Mendelian diseases to thousands of different human genes^1–3^. Thousands of additional rare Mendelian diseases and human genes await discovery. Frequency-based filters have proven extremely effective in providing diagnosis in such cases^4^. In essence, variants found in a control population (common variants) are likely to be benign^5^ while functional rare variants not found in the control population but seen in multiple affected individuals are likely to be disease causing^6–8^. These filters seek the gene or variant present in all (most) affected individuals but in no (very few) unaffected individuals.

For example, one can take a small cohort of unrelated individuals suspected of suffering from the same genetic disorder, and compare their genomes to that of tens of thousands of unaffected individuals (e.g., from the exome aggregation consortium, ExAC^5^). As we show below, in multiple scenarios, the gene with rare functional mutations in most patients in our small cohorts is indeed causal of their condition.

Frequency-based computation highlights the fundamental “serve or protect” dilemma of genomic data: **“Serve:”** to find the root cause of a patient’s disease, one wishes to compare a patient genome to as many other genomes as possible, both affected and unaffected, related and unrelated. Thus, to advance modern medicine, all sequenced genomes should be shared. **“Protect:”** one’s genome continues to reveal more and more about oneself, including susceptibility to a variety of diseases^9^. Sharing it with others can lead to discrimination and bias. To protect its owner and next of kin, no sequenced genome should be shared.

To date, this dilemma has been solved by allowing institutions unrestricted access to all the genomes in their possession. Limited sharing between institutions is done by providing obfuscated summary statistics^10^. Current commonly adopted methods for sharing have shortcomings that make them suboptimal. Providing full access at individual institutions allows for too much information to be shared in certain situations^11^. Disease-specific beacons are prone to attack and can end up identifying individuals participating in the study^12^. Beacons also only provide allele-presence query capabilities and do not have the flexibility needed for analyzing multifactorial variant interactions within an individual^13^. It is also risky to share genomic data in the clear with third-party services specializing in genomic and disease analysis. We are unaware of any cryptographically-secure method for sharing genomic data to perform computational operations that allow identifying causal variants in patients.

To better resolve this dilemma, we first note that while all of the genomic variants from all individuals are needed to perform the computation, only a handful of causal variants are ultimately of interest in the context of Mendelian patients (in the example above, just the rare variants in the single gene mutated in most patients).

We introduce here a modern, proof-of-concept cryptographic implementation which both serves and protects. The secure computation can be run on entire genomes (Serve), while no party involved in the computation learns anything about the inputs of the other participants except for the output which is computed together (Protect). We use real patient data to show that our secure implementation reveals minimal information while diagnosing patient genomes through 3 different strategies using practical amounts of compute time and memory. Cryptographic methods have been used in different genomic contexts such as microbiome analysis^14^, GWAS analysis^15^ and genomic alignment^16^, but this is the first implementation that we are aware of that is geared towards diagnosing Mendelian patients, a timely and potent need.

## Methods

### Representing genomic data as vectors

Assume each individual involved in a study has private access to their exome (or genome). If we are looking to identify a causal variant, we define a **variant vector** (long list of length 28,413,589) of all possible rare missense / nonsense variants in the human genome from the first gene on chromosome 1 to the last gene on chromosome Y. We provide a copy of this vector to each individual (affected and unaffected), and ask them to privately denote True/False next to each variant (to indicate whether they have the specific mutation or not, respectively). If we are looking to identify a causal gene, we provide each individual a **gene vector** of 20,663 genes in the human genome from A1BG to ZZZ3. We ask them to write “1” next to a gene if they have one or more rare functional variants in this gene, and otherwise, they write “0”. See Supplementary Figure 1A,B.

### Defining computations of interest (MAX, INTERSECTION, SETDIFF)

We define three operations used for patient diagnosis (Supplementary Figure 1C). Imagine two affected individuals are represented by two rare functional variant vectors (True/False lists). Intersecting these two vectors will reveal all the rare functional variants they share. Formally, we perform a Boolean **INTERSECTION** (or **AND**) operation (x AND y = True only if x = y = True, and otherwise, it is False) between all possible patient variants. Next, if we also have access to an unaffected family member, we can further exclude any variant they share. We do this with a Boolean set difference (**SETDIFF**) operation (x SETDIFF y = True only if x = True and y =False, otherwise it is False). Finally, imagine we have access to a small cohort of unrelated patients sharing a set of phenotypes. We would like to find the gene affected by one or more rare functional variants in the greatest number of patients within the cohort. For this, we use the patient gene vectors (0/1 lists). We sum 0/1s across patients for each gene, and then we use the maximum (**MAX**) operation to find the entry (gene) with the greatest number (of affected cases; Supplementary Figure 1C).

Remarkably, modern cryptography allows any number of individuals to jointly learn the final result of these MAX, SETDIFF, INTERSECTION operations without any of them learning anything else about each other’s genomes (or vectors).

### Encryption and decryption

An important cryptographic primitive we rely on is a secret-key encryption scheme. In a secret-key encryption scheme, a secret key is used to encrypt and decrypt messages with the guarantee that the encryptions of any two messages are indistinguishable, and yet, they can be successfully decrypted (to obtain the original message) given the key (Supplementary Figure 2).

### Secure multiparty computation

Multiple mathematical frameworks and computational implementations exist for secure multiparty computation on encrypted data. These provide different tradeoffs in complexity and efficiency^17^. In this work, we use Yao’s protocol to securely evaluate functions between two parties^18^. Abstractly, we write the function as *f*(*x*_1_, *x*_2_) where *x*_1_ denotes the input of the first party and *x*_2_ denotes the input of the second party. Any function *f*(*x*_1_, *x*_2_) can be represented by a combination of Boolean operations (for example, see Supplementary Figure 3). Yao’s protocol provides a way of evaluating the Boolean circuit (operator-by-operator) without revealing the inputs *x*_1_, *x*_2_. We illustrate this in detail in Figure 1.

**Figure 1.**
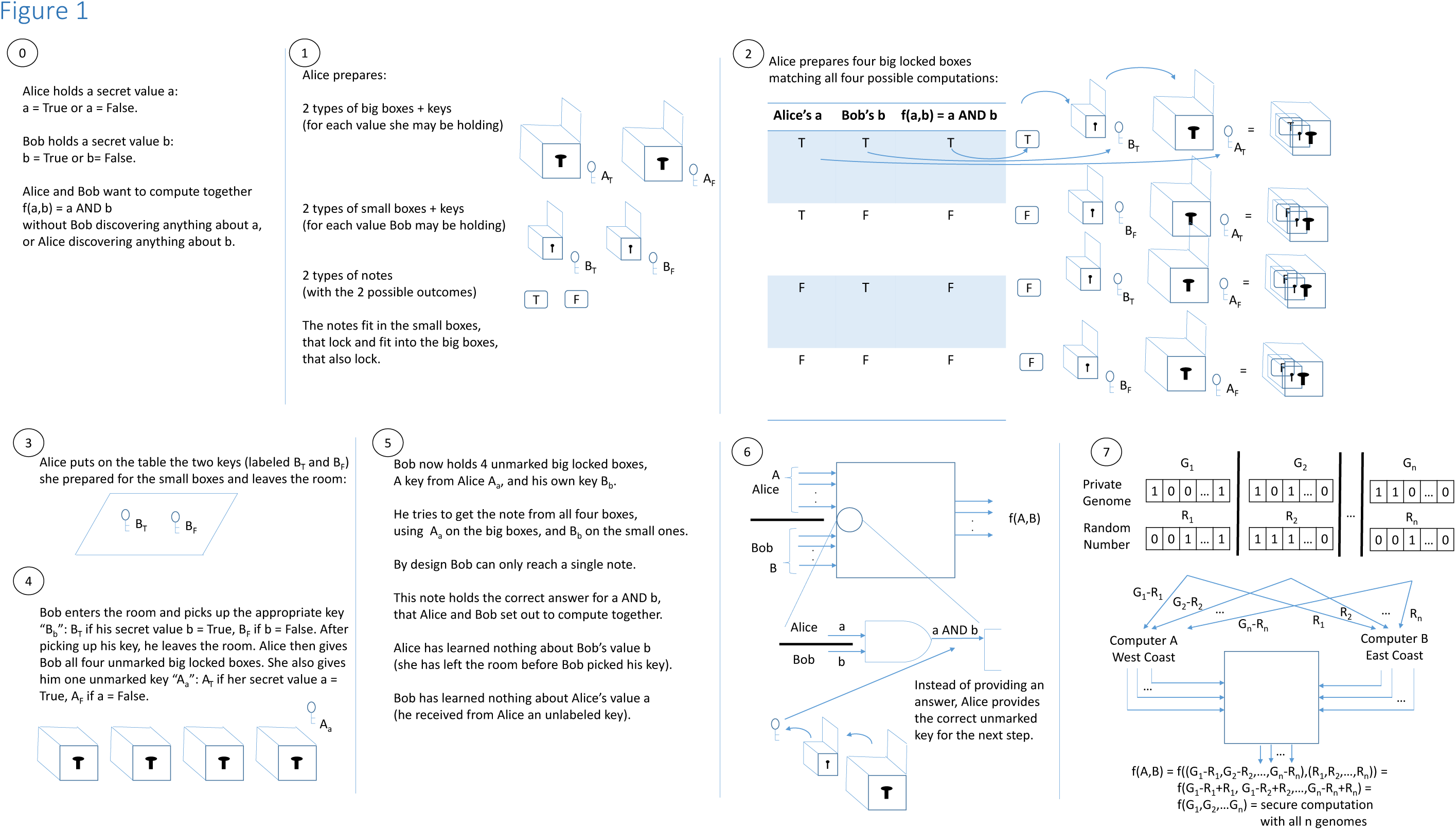
Yao’s protocol for secure multiparty computation. Steps 0-7 describe the overall secure protocol for computing any function F between two or more parties F(A, B, .. ,Y, Z). We first describe a secure two-party computation protocol between Alice (A) and Bob (B). **Step 0:** Alice and Bob are trying to compute a joint function without revealing their inputs to the other party. **Step 1:** Alice creates a key/box for each possible value for each input (0 or 1). **Step 2:** Alice double locks (double encrypts) each of the four possible outputs by placing the relevant output note in two boxes corresponding to each combination of the two inputs. **Step 3:** Alice gives Bob the option of choosing exactly one of two possible keys, labeled B_T_ and B_F_. **Step 4:** Bob picks up exactly one key B_b_ where b corresponds to his hidden input which only he knows (the oblivious transfer protocol ensures that Bob can only pick up *one* key). After Bob makes his selection, Alice shuffles the doubly-locked boxes and hands them to Bob along with the key A_a_ corresponding to her input a. Steps 1-4 is repeated for each of the inputs to the function. **Step 5:** For each operator in the function that depends only on input values (i.e., the first “layer” of the circuit), Bob has four doubly-locked boxes and two keys A_a_ and B_b_ but he does not know Alice’s input and Alice does not know Bob’s input. He uses A_a_ and B_b_ and tries to unlock all four boxes. Only one of the four doubly-locked boxes will successfully open, revealing the joint output without revealing Alice’s or Bob’s inputs. **Step 6:** The revealed output yields the key for the next operation (gate) in the circuit. Steps 5 and 6 are repeated for each operation in the function. At the end of the computation, instead of keys, Bob obtains the values that make up the output of the computation. **Step 7:** This secure two-party computation process can be expanded to N parties by using additive secret sharing between two non-colluding cloud servers. The N-input function is thus transformed into a two-input function.

While Yao’s protocol provides a simple and efficient solution for secure two-party computation, in many of the scenarios we describe, the computation occurs among multiple parties (e.g., many individuals, each with their personal genome). It is very straightforward to reduce the general problem of secure *multiparty* computation to that of secure two-party computation by working in the “two-cloud” model. In the two-cloud model, we assume that there are two *non-colluding* servers (e.g., these could be managed by two independent government agencies) that aggregate the inputs from each party in a privacy-preserving manner and then perform the computation. Each server on its own has *no knowledge* of the data as shown in Figure 1.7 (see Online Methods and Discussion).

### Protection quotient

We define the Protection Quotient as the fraction of private information that is not exposed (to neither the other participants nor the entity running the computation) during the computation. Using our encryption scheme, the Protection Quotient equals the total number of patient variants withheld from the output divided by the total number of patient variants input into the computation. Standard unencrypted patient diagnosis operations have a protection quotient of 0%, because all values must be exposed to perform the computation. All our applications below have a protection quotient of 97.1 - 99.7%, maximizing privacy while retaining full utility.

## Results

### Example Mendelian applications of secure computation

To prove the pragmatic utility of our approach, we demonstrate three different secure operations over real Mendelian patients where we successfully identify the causal variants in each scenario (Table 1):

**Table 1.** Summary of results for different secure genomic multiparty computation scenarios, all using real patient data.

| Operation | Relevant Information for each Operation |  |  |  |  |  |  | Running Time Measurements |  |  |
| --- | --- | --- | --- | --- | --- | --- | --- | --- | --- | --- |
| (a) MAX<br>(over genes) | Scenario:<br>Small disease cohort | # unrelated probands (who avoid openly sharing their data) | Rare functional variants (genes) per proband (median) | # probands with rare functional variant/s in gene (top 3, descending order) | Gene name | Proven causal gene for disease | Protection quotient (1- # of variants shared of top gene/total # of variants) | Bandwidth (GB) | Compute (sec) | Network (sec) |
|  | Freeman Sheldon Syndrome | 3 | 258 (253) | 3 | MYH3 | MYH3 | 1 - 3/767 = 99.6% | 0.02 | .15 | 4.91 |
|  |  |  |  | 2 | DBT |  |  |  |  |  |
|  |  |  |  | 1 | ACADVL |  |  |  |  |  |
|  | Hajdu-Cheney Syndrome | 7 | 278 (272) | 6 | NOTCH2 | NOTCH2 | 1 - 8/1853 = 99.6% | 0.03 | .18 | 7.29 |
|  |  |  |  | 3 | HLA-DRBI |  |  |  |  |  |
|  |  |  |  | 3 | MCC |  |  |  |  |  |
|  | Kabuki Syndrome | 10 | 262 (257) | 8 | KMT2D | KMT2D | 1 – 8/2754 = 99.7% | 0.04 | .22 | 9.59 |
|  |  |  |  | 3 | COL6A1 |  |  |  |  |  |
|  |  |  |  | 3 | FLNB |  |  |  |  |  |
|  | Miller Syndrome | 4 | 267 (258) | 4 | DHODH | DHODH | 1 – 8/1063 = 99.3% | 0.03 | .18 | 7.29 |
|  |  |  |  | 3 | DNAH5 |  |  |  |  |  |
|  |  |  |  | 2 | ACOX2 |  |  |  |  |  |
| (b) SETDIFF<br>(over variants) | Scenario: familial | Patient ID (avoid sharing w provider) | # rare functional variants | # proband only variants (revealed) | Gene name | Proven causal gene | Protection quotient | Bandwidth (GB) | Compute (min) | Network (min) |
|  | Trio | 115-f | 185 | N/R | N/R | ACTB | 1 - 2/524 = 99.6% | 18.1 | 1.7 | 56.7 |
|  |  | 115-m | 164 | N/R | N/R |  |  |  |  |  |
|  |  | 115-a1 | 175 | 2 | ACTB |  |  |  |  |  |
|  |  |  |  |  | USH2A |  |  |  |  |  |
| (c) INTERSECTION<br>(over variants) | Scenario: 2 Hospitals | # suspicious variants (not shared) | Total intersecting variants (for patient phenotype comparison follow-up) |  |  |  | Protection quotient | Bandwidth (GB) | Compute (min) | Network (min) |
|  | Washington | 5,734 | 159 |  |  |  | 1 – 318/11,067 = 97.1% | 3.1 | 0.37 | 9.4 |
|  | Baylor | 5,333 |  |  |  |  |  |  |  |  |

### MAX identifies the causal gene in small patient cohorts with protection quotient above 99%

We use 4 small cohorts of unrelated individuals, suffering from very different rare diseases: Freeman Sheldon Syndrome (FSS), Hadju-Cheney Syndrome (HCS), Kabuki Syndrome (KaS) and Miller Syndrome (MiS). Each individual holds a private list of 211-374 rare functional variants in 210-356 genes (total 767-2,754 variants per computation). We use the secure MAX function to reveal only the top gene mutated across patients in each cohort. In all 4 cohorts, we find that the gene mutated in most individuals is the one that has been proven to be the causal gene: MYH3 in FSS^6^, NOTCH2 in HCS^19^, KMT2D in KaS^8^ and DHODH in MiS^7^ (Table 1a).

Secure computation only reveals the variants in the most mutated gene in each cohort while protecting the remaining 764 variants in FSS, 1845 variants in HCS, 2746 variants in KaS and 1055 variants in MiS. This computation has a protection quotient of 99.3-99.7% for all 4 cohort disease datasets. The computation is performed over all 20,663 genes and completes in just 5 - 10 seconds, with one server on the East Coast and the other on the West Coast (Table 1a). The total protocol execution time, bandwidth and compute time all grow logarithmically with the number of cohort individuals involved in the secure computation (Supplementary Figure 4A).

### SETDIFF identifies the causal variant in a trio with protection quotient 99.6%

Unaffected mother and father, and affected male child with female external genitalia, each holds a list of 164-185 (total 524) rare functional variants found in their exomes. The secure SETDIFF operation reveals to the family and test providers only 2 rare variants found in the child but in neither parent (Table 1b). Literature review provides a diagnosis based on one of these two variants: the ACTB gene^20^.

Secure computation keeps 522 variants private while sharing only 2 variants with the test provider and all individuals involved in the computation. This computation has a protection quotient of 99.6%. Because three parties are now involved, the total computation time using a single server thread on either coast is 57 minutes (Table 1b). However, the variant list can easily be split between a small computer array on either coast, such that a typical 30-node cluster brings computation time down to under 2 minutes. The protocol execution time, bandwidth and compute time all grow logarithmically with the number of family members involved in the secure computation (Supplementary Figure 4B).

### INTERSECTION identifies patients of interest across 2 hospitals with protection quotient 97.1%

Two or more genome centers may want to compare their patient lists to see if together they can find multiple patients with the same rare functional mutation, and similar phenotypes, while revealing nothing else to each other. For example we took 928 Washington Mendelian Center (WMC) patients and 282 Baylor Hopkins Center (BHC) patients. For each hospital we prepared a list of over 5,000 rare functional variants seen in one or more of their patients. Using the secure AND function, the two hospitals find a short list of just 159 variants present in both hospitals, pointing at patients who would benefit from phenotype comparison. This short list includes “positive controls” such as known disease variant NOTCH1:p.E694K, associated with partial/incomplete penetrance of aortic valve disease^21^. Indeed the WMC and BHC patients are phenotypically characterized with left ventricular outflow defect and thoracic aortic aneurysm, respectively. The list also offers exciting novel gene-disease associations such as rare functional variant HCN3:p.R648H (with frequency 5.47·10^-5^ and 0 in ExAC and 1000 genomes data, respectively). HCN3 is a voltage-gated cation channel gene, whose mouse knockout causes abnormal ventricular action potential waveform^22^. Promisingly, in patients from WMC and BHC, this mutation is correlated with dilated cardiomyopathy and coarctation of the aorta, respectively.

Secure computation only reveals 2x159 potential causative variants while protecting the remaining 10,749 variants with a protection quotient of 97.1%. This computation is performed over all rare functional variants in the exome with a total protocol execution time of 9.4 minutes using a single server thread on either coast (Table 1c). Because every variant is evaluated independently, a 30-node compute cluster on either end will reduce total computation time to below 20 seconds. As we learn to appreciate Mendelian mutations outside of the exome, the total time, bandwidth and compute time scale linearly with the size of the variant list shared for secure computation (Supplementary Figure 4C).

## Discussion

Rare diseases are cumulatively common (some estimate that 10% of the US population are affected with rare disorders). About 7,000 rare Mendelian conditions have been described to date. Of these, approximately 4,000 have been definitively diagnosed as single gene diseases, mapping to over 4,000 genes in the genome. The procedures we describe are applicable for all of these. There are only a handful of medical conditions diagnosed with certainty to the interactions of just 2 genes. Far less is known for diseases caused by more than 2 genes. Personal Genomics poses a fundamental “serve or protect” dilemma: should one serve their genome in the service of better diagnosis and ultimately disease eradication, or should one protect oneself and next of kin against potential discrimination by refusing to share their genome. This dilemma is particularly evident in the field of Mendelian diseases. It is essential to develop tools and methods to effectively share genomes while maintaining their privacy and security. Because genome privacy is best served where a definitive diagnosis exists, we focus on single disease gene discovery and diagnosis. Here we present a secure approach for multiple parties to perform exact computations that diagnose Mendelian diseases, while keeping all participating genomes private.

The scenarios we present are all real. Genome privacy is extremely appealing in all of them: Complete strangers in disease cohorts (Table 1A) learn nothing about each other except their shared disease-causing gene mutations. For participants where the assay does not provide an answer, absolutely nothing is revealed. In larger family trees, more distantly related members will appreciate genome privacy. Even in a young nuclear family (e.g., a trio; Table 1B), the test provider learns almost nothing except the likely disease-causing mutation in the offspring. Moreover, they learn virtually nothing about the parents themselves. In the two hospital scenario (Table 1C), only variants that are worthwhile comparing are revealed while the vast majority of variants remain private to each institute’s researchers and patients.

In all of these cases, the quantities revealed and those that remain private are a privacy advocate’s dream come true: We predominantly reveal only variant/s crucial for patient diagnosis, family counseling and any potential treatment. What remains private is predominantly variants of unknown significance (VUS) that are of little value for diagnosing one’s medical condition. However, these same VUS variants almost certainly uniquely identify a person as a participant in an analysis, and have the potential to reveal now or in the future other personal traits that may be further cause for discrimination.

For this proof-of-concept work, we assume that the protocol participants are “honest-but-curious,” (sometimes referred to as “semi-honest”)—that is, we assume that the parties are properly incentivized to *honestly* follow the protocol, but at the end of the protocol execution, they may try to learn some additional information (about other parties’ inputs) based on the messages they receive during the protocol execution. We say that a protocol is secure if the only information any party learns by participating in the protocol can be inferred just from that parties’ input and the overall output of the computation. In other words, none of the parties should be able to learn something about another parties’ input other than what is explicitly revealed by the output of the function.

Yao’s protocol gives an efficient solution for secure two-party computation in the presence of semi-honest adversaries^23^. We note that there are well-established ways to extend Yao’s protocol to additionally provide security against malicious parties who deviate from the protocol description in order to compromise the privacy of other participants or corrupt the results of the computation^24^. In addition, protecting against participants that submit malicious (or malformed) inputs to the protocol can be done by ensuring that if a participant’s variant vector does not meet certain criteria, or is not accompanied by an appropriate certificate, then the computation aborts and does not produce any output. Furthermore, in this paper, we introduce an operation-specific “protection quotient”, a novel metric to assess the fraction of information secured by the computation. The protection quotient can be used to further restrict the output returned to all parties if the defined privacy requirements are not met. For instance, if a trio analysis results in more than a few expected de novo exome mutations, only an error message will be produced. This approach is preferred for example to differential privacy^25,26^ which adds random genomic variation as noise into aggregated summary statistics to try and avoid individual identification in pooled genomics data^15^.

The basic principle underlying our design is to perform exact secure computation on the complete (private) genomes of all participating individuals. This is in direct contrast to the more traditional and less effective routes of publishing obfuscated frequencies aggregated across multiple individuals. The computational resources we use to retain genomic privacy are not negligible, yet are perfectly within the capabilities of off-the-shelf modern computers to complete the operation in seconds or minutes, even when communicating between the East and West coasts. And while no security mechanism may be perfectly impenetrable, it is certainly preferable to have a security mechanism in place (especially if it allows for *exact computation*) where none currently exist. Many further extensions and applications of our computational framework are possible, and are sure to provide incentives for the development of more secure and faster methods. A widespread deployment of computer libraries efficiently implementing these principles will encourage individuals to securely contribute their genomes for the common good, and thus greatly fuel advances in both personal genomics and privacy in the 21^st^ century.

## On-Line Methods

### Patient datasets

Whole exome sequences of patients were obtained from dbGaP studies phs000204.v1.p1^6^ (Freeman Sheldon Syndrome), phs000244.v1.p1^7^ (Miller Syndrome), phs000295.v1.p1^8^ (Kabuki Syndrome), and phs000477.v1.p1 (Hajdu-Cheney Syndrome). Pre-processed variant call format (VCF) files for patients from 2 Centers for Mendelian Genomics were obtained from dbGaP studies phs000693.v4.p1 (University of Washington), and phs000711.v3.p1 (Baylor Hopkins). Our trio family was obtained from Stanford Hospital. All human subject research was performed under guidelines approved by the Stanford Institutional Review Board.

Sequencing reads were mapped to the GRCh37/hg19 assembly of the human genome using BWA MEM v0.7.10-r789^27^. Variants were called using GATK v3.4-46-gbc02625 following the HaplotypeCaller workflow from the GATK Best Practices^28^.

### Variant annotation

ANNOVAR v527 was used to annotate variants with predicted effect on protein coding genes using gene isoforms from the ENSEMBL gene set version 75 for the hg19/GRCh37 assembly of the human genome^29,30^. All canonical gene isoforms were used where the transcript start and end are marked as complete and the coding span is a multiple of three.

### Cryptographic techniques

In a secure multiparty computation (MPC) protocol^18,31^, a group of users (often called *parties*) seek to jointly compute a function over their inputs without revealing any additional information about their particular inputs. The function that the parties compute is determined based on the specific scenario. The computation consists of several rounds of interaction, where in each round, the parties exchange a series of messages. At the conclusion of the protocol, each participant learns the output of the computation evaluated on everyone’s joint input. No additional information beyond the explicit output is revealed to any party (the process is abstracted in Figure 1).

Every arithmetic computation can be expressed as a sequence of Boolean logical operations (that is, operations on bits {0,1}). This is precisely how the modern computer works. Yao’s protocol allows two users, Alice and Bob, to compute arbitrary functions over their inputs. More precisely, if Alice has an input *x* and Bob has an input *y*, Yao’s protocol allows them to compute *f*(*x*, *y*) in a way such that Alice learns nothing about *y* and Bob learns nothing about *x* other than the output value *f*(*x*, *y*). In general, expressing a function in terms of Boolean operations greatly increases the computational cost of evaluating the function. To maximize the efficiency of Yao’s protocol, it is important to choose functionalities with simple or compact representations as Boolean circuits. An example of a Boolean circuit is shown in Supplementary Figure 3.

In this work, we cast diagnosing Mendelian patients as (simple) arithmetic/logic computations that admit efficient Boolean circuit representations. We now describe how the secure computation protocols work. To do this we first introduce two standard tools from cryptography: (1) symmetric (secret-key) encryption^32^ and (2) oblivious transfer^33–35^.

### Encryption and decryption

A secret-key encryption scheme consists of two functions: **Encrypt and Decrypt**. The encryption function takes a cryptographic key *k* and a message *m* and outputs a ciphertext *c*. The decryption function takes the cryptographic key *k* and a ciphertext *c* and outputs a message *m*. Intuitively, encryption and decryption are inverse operations: if we encrypt a message under a key *k*, decrypting the resulting ciphertext with the *same* key *k* recovers the original message. More precisely, we can say that for any key *k* and any message *m*, Decrypt(*k*, Encrypt(*k*, *m*)) = *m*. In a *symmetric* (or secret-key) encryption scheme, both the encryption and the decryption functions require knowledge of the secret cryptographic key. The key is a random string drawn from some key-space. The precise nature of the key-space varies depending on the details of the encryption scheme, and is immaterial to our presentation in this paper. An encryption scheme is considered to be secure if the ciphertext does not reveal any information about the underlying message to any user who does not possess the secret encryption key (certainly, a user who holds the secret key can decrypt and learn the message). One way to formalize this is to say that a user who does not have the encryption key is unable to tell an encryption of a message *m*_0_ apart from an encryption of another message *m*_1_. In other words, ciphertexts hide all information about their underlying message to all users who do not have the encryption key. We illustrate this in Supplementary Figure 2.

Under this definition, messages can also be encrypted multiple times. For instance, a message *m* can be “double encrypted” under two keys *k*_1_ and *k*_2_ by first encrypting *m* using *k*_1_ and then encrypting the resulting ciphertext using the second key *k*_2_. This procedure yields another ciphertext. Decryption proceeds by first decrypting with key *k*_2_, and then decrypting the result (a ciphertext) with *k*_1_. In particular, we can write

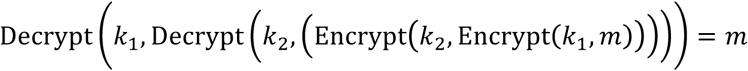

Security of the double encryption scheme follows directly from the security of the underlying encryption scheme. In particular, a user who does not have both *k*_1_ and *k*_2_ cannot learn any information about the underlying message that has been doubly encrypted using *k*_1_ and *k*_2_. Numerous symmetric (secret-key) encryption schemes exist in the literature^32^.

### Oblivious transfer

An oblivious transfer (OT) protocol^33–35^ is a two-party protocol between a sender and a receiver. An OT protocol enables the receiver to selectively obtain one of two possible messages from the sender without revealing to the sender which message the receiver requested. More precisely, the sender holds two messages, denoted *k*_0_ and *k*_1_ and the receiver holds a selection bit *b* ∈ {0,1}. At the end of the OT protocol, the receiver obtains the chosen message *k*_*b*_ and learns nothing about the other message *k*_1–*b*_. The sender does not learn anything about the receiver’s choice bit *b*. Numerous oblivious transfer protocols have been proposed in the literature^33–35^.

### Overview of steps for secure computation

In a secure two-party computation protocol, Alice holds an input *x* ∈ {0,1}^*n*^ and Bob holds an input *y* ∈ {0,1}^*n*^. We write {0,1}^*n*^ to denote a binary input of length *n* (e.g., for instance, *n* could be of the binary representation of the variant vector or the gene vector we define in our main text). Their goal is to compute a function *f*(*x*, *y*) on their joint input (*x*, *y*). The computation is considered “secure” if at the end of the computation, the only information that Alice and Bob learn is the function value *f*(*x*, *y*) and nothing else about the other party’s input. It is important to note here that the function output *f*(*x*, *y*) could reveal some information about the inputs *x* and *y* (for example, in our trio scenario, whatever de novo variant we report in the child, we can deduce by definition does not exist in either parent). As we note in the Discussion section, we work in the honest-but-curious model where we assume that Alice and Bob follow the protocol specification as directed, but may, at the end of the protocol execution, try to infer some additional information about each other’s private input. We now describe how Yao’s protocol can be used to securely evaluate any function over two inputs in the honest-but-curious model.

To apply Yao’s protocol, it is first necessary to represent the function *f* as a Boolean circuit on inputs *x* and *y*. At the most basic level, the building blocks we have are AND (x AND y = True only if x = y = True, otherwise it is False) and XOR (exclusive-or, x XOR y = True only if x = True and y = False, or if x = False and y = True, otherwise it is False) gates. These basic gates can be combined to obtain circuits of arbitrary expressive functionalities^17^. In our description below, we will oftentimes refer to the concrete example of securely evaluating the AND function on single-bit inputs (above). A visualization of the complete protocol is given in Figure 1.

### Overview of Yao’s secure two party computation protocol

We now describe Yao’s protocol. For ease of presentation, we present a simplified (but less efficient) description of Yao’s protocol here. Our implementation (based on the JustGarble^36^ library) follows the high-level blueprint described here, but includes several optimizations, notably the free-XOR^37^ and half-gate^38^ optimizations.

#### Step 1

For each wire in the circuit, Alice chooses two keys. Recall that in a Boolean circuit, each wire in the circuit can take on two possible values (0 or 1; sometimes also referred to as False and True, respectively). Alice associates one of the keys with the wire value 0 and another key with the wire 1. For the particular case of securely evaluating the AND function *z* = *x* ∧ *y*, Alice picks three pairs of keys and associates one pair with each of *x*, *y*, and *z*. We denote these keys 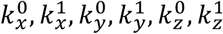. In this example, 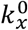 is the key associated with the input bit *x* taking on the value 0 and 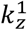 is the key associated with the output bit *z* taking on the value 1. This step is shown in Figure 1.1.

#### Step 2

For each gate in the circuit, Alice constructs a “garbled” truth table. For each row in the truth table, the algorithm takes the key associated with the value of the output wire and *double* encrypts it using the two keys associated with the values of the two input wires. For the particular case of evaluating a single AND gate, Alice would construct the following table of ciphertexts

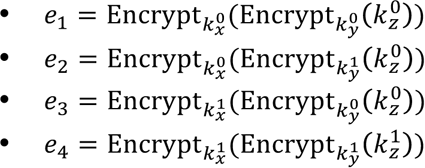

For the output wires of the circuit, instead of double encrypting an encryption key, Alice directly double encrypts the value of the output wire (e.g., 0 or 1). This step is shown in Figure 1.2.

#### Step 3

After garbling the circuit (Steps 1-2 / Figure 1.1-1.2), the secure computation begins with Bob using the oblivious transfer protocol (above) to obtain the keys for the input wires associated with his input. The oblivious transfer protocol ensures the following: Bob only learns one of the two keys associated with each of his input wires (this will ensure that Bob can only evaluate the function on a single set of inputs), and Alice does not learn which wire Bob requested (that is, Alice does not learn Bob’s input). For the particular case of evaluating a single AND gate, if Bob’s input is *b* ∈ {0,1}, then Bob would play the role of the receiver in an OT protocol with input *b*. Alice would play the role of the sender with messages, 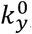, 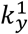 (the keys associated with Bob’s input wire). At the end of the oblivious transfer protocol, Bob obtains 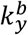 (the key associated with his input), and learns nothing about the key associated with the complement of his input (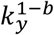). Alice learns nothing about Bob’s input *b*. This step is shown in Figure 1.3.

#### Step 4

After Bob receives the keys associated with his input via the oblivious transfer protocol, Alice sends Bob the garbled tables associated with each gate (after randomly permuting the rows of each table). Additionally, Alice sends Bob the wire encodings of her input. For the particular case of evaluating a single AND gate, if Alice’s input is *a* ∈ {0,1}, Alice would send 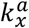 (the key associated with *x* = *a*) to Bob. This step is also shown in Figure 1.4.

#### Step 5

With all this information, Bob can complete the function evaluation and compute the output. In particular, after Steps 3 and 4, Bob should have a single key for each of the input wires of the Boolean circuit. Then, for each gate in the circuit, Bob takes the input keys he has and attempts to decrypt the rows in the garbled table associated with that gate. Because the entries in the garbled table are *double encrypted* using the keys associated with the input wires to the gate and Bob only has a *single* key for each of the wires, Bob is only able to decrypt a *single* row in the garbled table as shown in Figure 1.5.

#### Step 6

In doing so, Bob is able to learn one of the keys associated with the gate’s output wire (moreover, by construction of the garbled table, the output key Bob obtains is precisely the one associated with the value corresponding to evaluation of the gate on the input bits). Thus, starting with the input wires, Bob is able to evaluate the circuit gate-by-gate as seen in Figure 1.6. Once Bob reaches the output layer of the circuit, he is able to decrypt the ciphertexts and obtain the value of each output wire.

**Summary.** To summarize, in Yao’s secure two-party computation protocol, Alice begins by constructing a garbled truth table for each gate in the Boolean circuit. She does so by double encrypting each row in the truth table (using the keys associated with the input bits). She gives Bob the garbled truth tables as well as the keys associated with her input. Using oblivious transfer, Bob obtains the keys associated with his input. Armed with a single key for each of the input wires in the circuit, Bob is able to evaluate the garbled circuit gate-by-gate. For each gate, Bob takes his input keys and uses them to decrypt one of the rows of the garbled table associated with the gate. This yields the key associated with the particular wire. Finally, at the end of the computation, Bob decrypts the ciphertexts associated with the output wires to learn the output of the circuit. Bob then sends the result of the computation to Alice.

### Extending Yao's protocol to N parties

Yao’s protocol allows two parties to securely evaluate an arbitrary function. However, in general, we desire to compute across a large number of parties (e.g., study participants). While there are secure multiparty computation protocols that support more than two parties, (e.g., the SPDZ^39^, GMW^31^, or BGW^40^ protocols), a key limitation of these protocols is that they require all participating parties to be online during the protocol execution. Moreover, the number of rounds of communication in the protocol often grows with the complexity of the computation (note that this is in direct contrast with Yao’s protocol which is a two-round protocol, regardless of how complicated the computation is). As a result, there are substantial engineering hurdles to deploying these general protocols for multiparty computation across a large number of parties. In some cases (e.g., BGW^40^ and GMW^31^), the total bandwidth also scales quadratically in the number of parties, further limiting the practicality of these protocols.

A more efficient solution for general multiparty computation that avoids both the requirement that participating parties be online during the protocol execution as well as the potential communication blowup is to work in a “two-cloud” model. In this model, we assume there are two *non-colluding* cloud servers that facilitate the protocol execution. At the beginning of the protocol execution, each of the participating parties “split” their inputs and share it with the two cloud servers. As long as the two clouds do not collude with each other, they do not learn anything about the inputs to the computation. After the two cloud servers have received the inputs from each of the participating parties, they engage in a two-party secure computation protocol (such as Yao’s protocol) to compute the function of interest. Notably, the parties that contributed the data do not have to be online during this step of the protocol. And moreover, communication is only necessary between the parties and the cloud servers; parties in particular do not have to communicate with each other. In a practical deployment, these two cloud servers might be managed by distinct governmental organizations within the NIH or WHO. Thus, by working in the two-cloud model, it is possible to transform any computation between *n* individuals into a secure two-party computation between two non-colluding parties.

#### Step 7

We now describe how to secure evaluate any functionality in the two-cloud model. Suppose there are *n* parties participating in the protocol execution and let *x*_1_, …, *x*_*n*_ denote their private inputs. To secure compute a function *f*, each of the *n* participants chooses a random value *r_i_* and sends *r_i_* to one of the two cloud servers. They then send to the other cloud server the value *x_i_* − *r_i_* note that the subtraction is performed modulo a large integer *N*). Once every party has submitted their inputs *r_i_* and *x_i_* − *r_i_* to the two cloud servers, the first cloud server has a vector of random values *a* = (*r*_1_, …, *r_n_*) and the second cloud server has a vector of random differences *b* = (*x*_1_ − *r*_1_, …, *x_n_* − *r_n_*). Because the subtraction is taking place modulo *N*, the values in *b* are distributed uniformly and more importantly, independently of the *x_i_*’s. The pair (*r_i_*, *x_i_* − *r_i_*) is often referred to as an “additive secret sharing” of the input *x_i_*. The property that this additive secret sharing scheme satisfies is that a single share reveals no information about the input, but two shares completely define the input. This means that as long as the two cloud servers do not collude, they learn no information about each party’s input (since they each possess just one share of the secret).

To complete the secure computation (of a function *f*), the two cloud servers simply apply Yao’s protocol to the following two-party functionality:

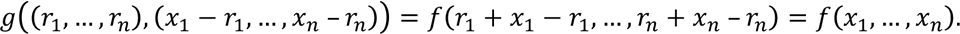

In other words, the two clouds compute the functionality that takes as input two vectors (each containing *n* values) and outputs the function *f* evaluated on the components corresponding to the sum of the two input vectors. Since summing the input vectors in this case reconstructs each party’s input, this procedure corresponds precisely to evaluating *f* on the parties’ inputs. Moreover, the two cloud servers do not learn any additional information about any particular party’s input because the evaluation of *g* is performed using Yao’s protocol (which is a secure two-party computation protocol). This procedure is shown in Figure 1.7.

### Constructing our Boolean circuits

As described above, arbitrary Boolean circuits can be constructed using only AND and XOR gates. To efficiently represent our set-intersection-based algorithms as Boolean circuits, we first construct some intermediate building blocks from the basic AND and XOR gates. The intermediate building blocks we require include addition circuits, comparison circuits, and equality circuits. For these building blocks, we use the circuits by Kolesnikov et al.^41^ (see Supplementary Figure 5).

### Software implementation

In our implementation, we use the JustGarble library^36^ for our implementation of Yao’s garbled circuits, and we use the Asharov et al.^42^ implementation of the oblivious transfer protocols. For better performance, we also implement the half-gates optimization^38^ for Yao’s garbled circuits. This implementation will be released upon publication. For our benchmarks, we set up a client and server on Amazon EC2 (to simulate the two cloud providers), and measure the total compute time, bandwidth, and overall protocol execution time (taking into account the network communication). We run our experiments on two memory-optimized EC2 instances (M4.2xlarge). Each instance runs an 8-core 2.4 GHz Intel Xeon E5-2676 v3 (Haswell) processor and has 32 GB of memory. While our protocols are naturally parallelizable, we use a single thread of execution in all of our experiments, and do not take advantage of the available parallelism. To simulate the non-colluding two cloud model, we used a wide-area network (WAN) setting where the two servers are far apart. We placed one of the servers on the West Coast (specifically, in the Northern California availability zone) and the other on the East Coast (specifically, in the Northern Virginia availability zone).

## Author Contributions

KJ, DW, DB and GB designed the study, analyzed results and wrote the manuscript. KJ and JB processed patient data. KJ and DW wrote software for the analysis.

## Acknowledgements

We thank Dr. Jon Bernstein and members of the Boneh and Bejerano labs for valuable discussions and project feedback. We also thank Stanford patients and clinicians, as well as the patients and professionals involved in the deposition of the dbGaP sets we use.

## Supplementary Figures and Tables

**Supplementary Figure 1.**
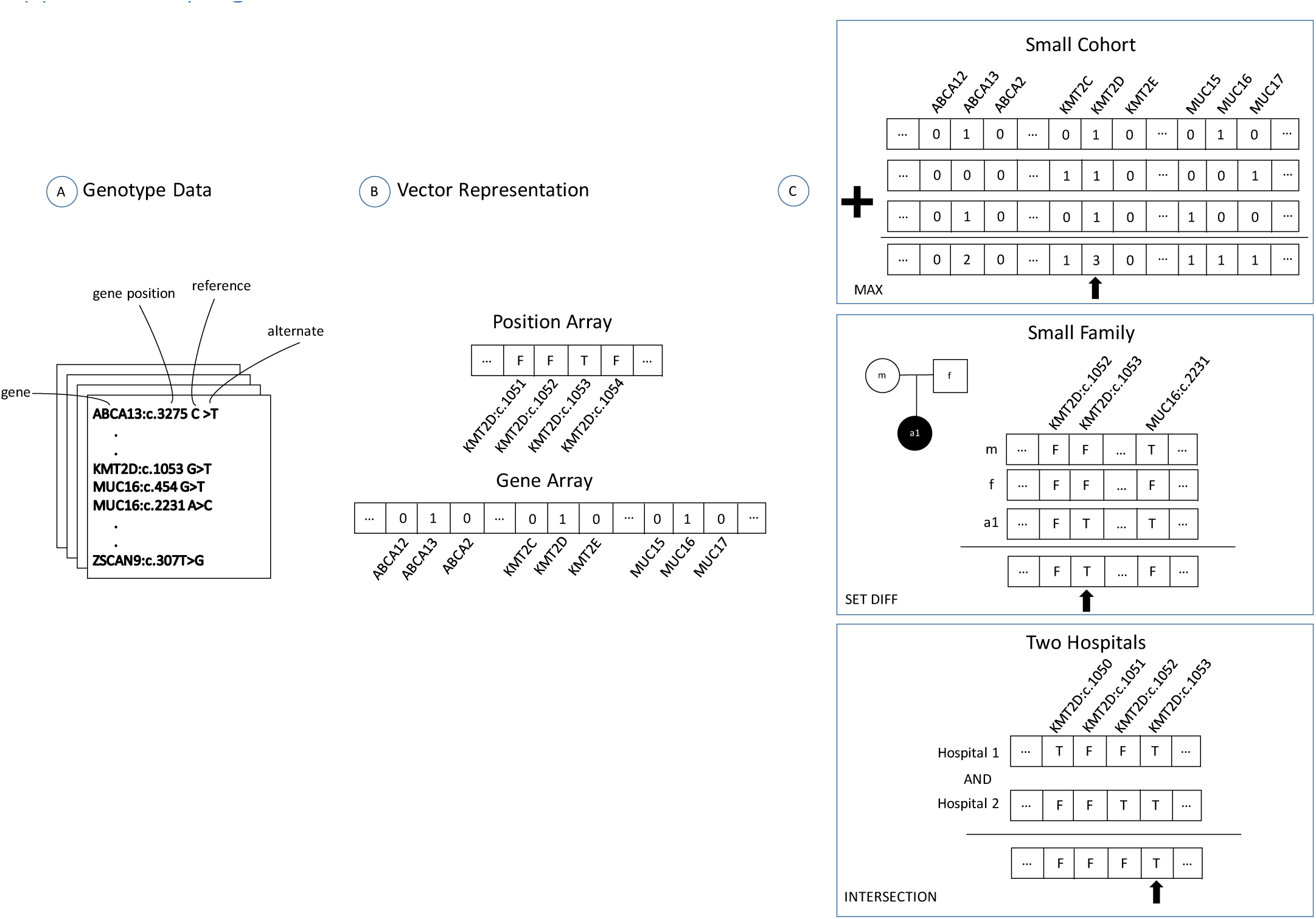
Representing genomic data as vectors for secure computation. (A) Each individual holds their personal genome private. (B) They are asked to fill in a position array/vector with True and False values depending on whether they have a rare functional variant at the listed position, or a 0/1 value in a gene array depending on whether they have none/some rare functional variant/s in each listed gene. (C) The resulting position/gene vectors are used to obtain the results of Table 1.

**Supplementary Figure 2.**
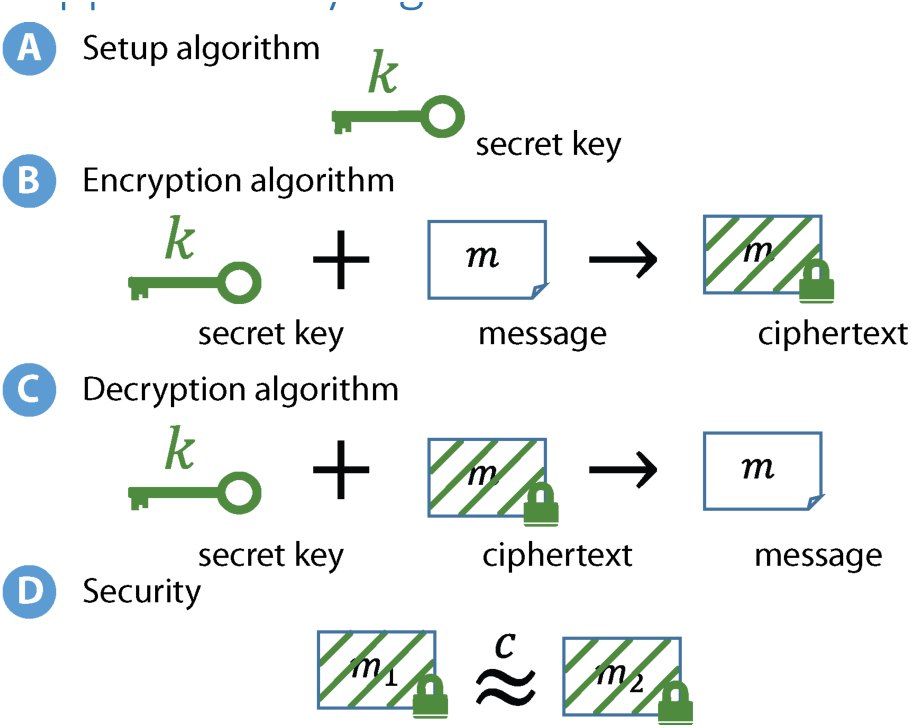
Encryption and decryption overview. A secret-key encryption scheme consists of three algorithms: (A) a setup algorithm which outputs a secret key (usually a long random string); (B) an encryption algorithm that takes in a secret key *k* and a message *m* and produces an encryption of *m* (called a ciphertext); and (C) a decryption algorithm that takes in the same secret key *k* and a ciphertext and produces the original message. We write Enc_*k*_(*m*) to denote an encryption of the message *m* under the secret key *k*. The correctness requirement for an encryption scheme states that decrypting the ciphertext output by Enc_*k*_(*m*) using the secret key *k* should yield the original message (plaintext) *m*. (D) The security requirement for a secret-key encryption scheme states that anyone who does *not* possess the secret-key *k* cannot distinguish an encryption of a message *m*_1_ from an encryption of a message *m*_2_, *irrespective* of the choice of messages *m*_1_ and *m*_2_. In other words,without the secret key, the ciphertext does not reveal any information about the encrypted message.

**Supplementary Figure 3.**
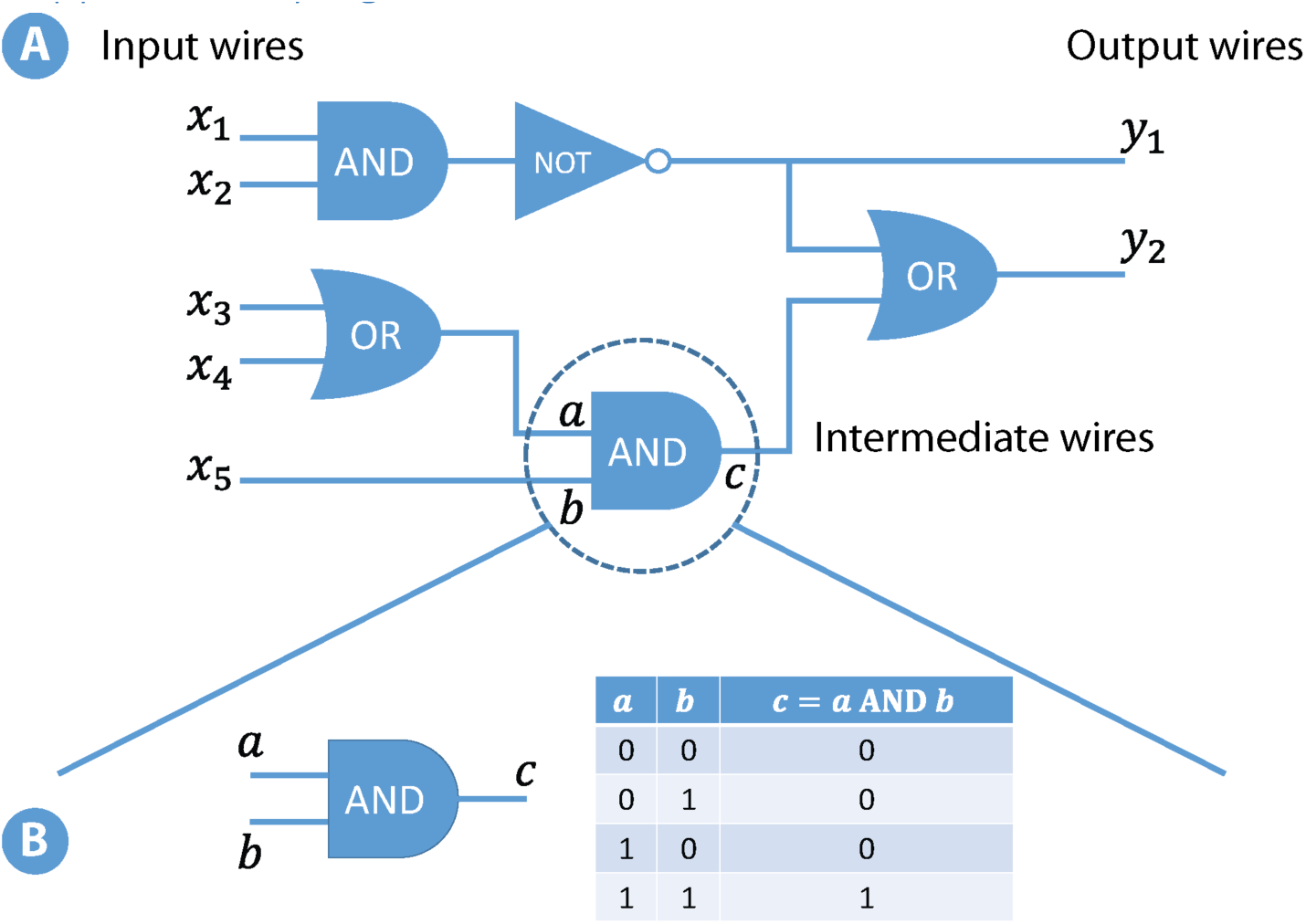
Computation using circuits. (A) A Boolean circuit consists of a sequence of logic gates (e.g., AND gates, OR gates, and NOT gates). Each logic gate takes one or two bits as input and produces a single bit of output. In a Boolean circuit, the outputs of one logic gate can be used as the input to another logic gate. We refer to these values as the *intermediate* values in the computation. In the circuit depicted in the figure, the inputs to the circuit are denoted *x*_1_, *x*_2_, *x*_3_, *x*_4_, *x*_5_ and the outputs of the circuit are denoted *y*_1_, *y*_2_. Specifically, this particular circuit implements a function over five input bits and produces two output bits. (B) Each gate in the Boolean circuit can be described by a truth table that specifies the mapping between each configuration of the input bits to a corresponding output bit. In the case of an AND gate, there are two input bits, and the output is 1 if and only if both input bits are 1. Otherwise, the output is 0.

**Supplementary Figure 4.**
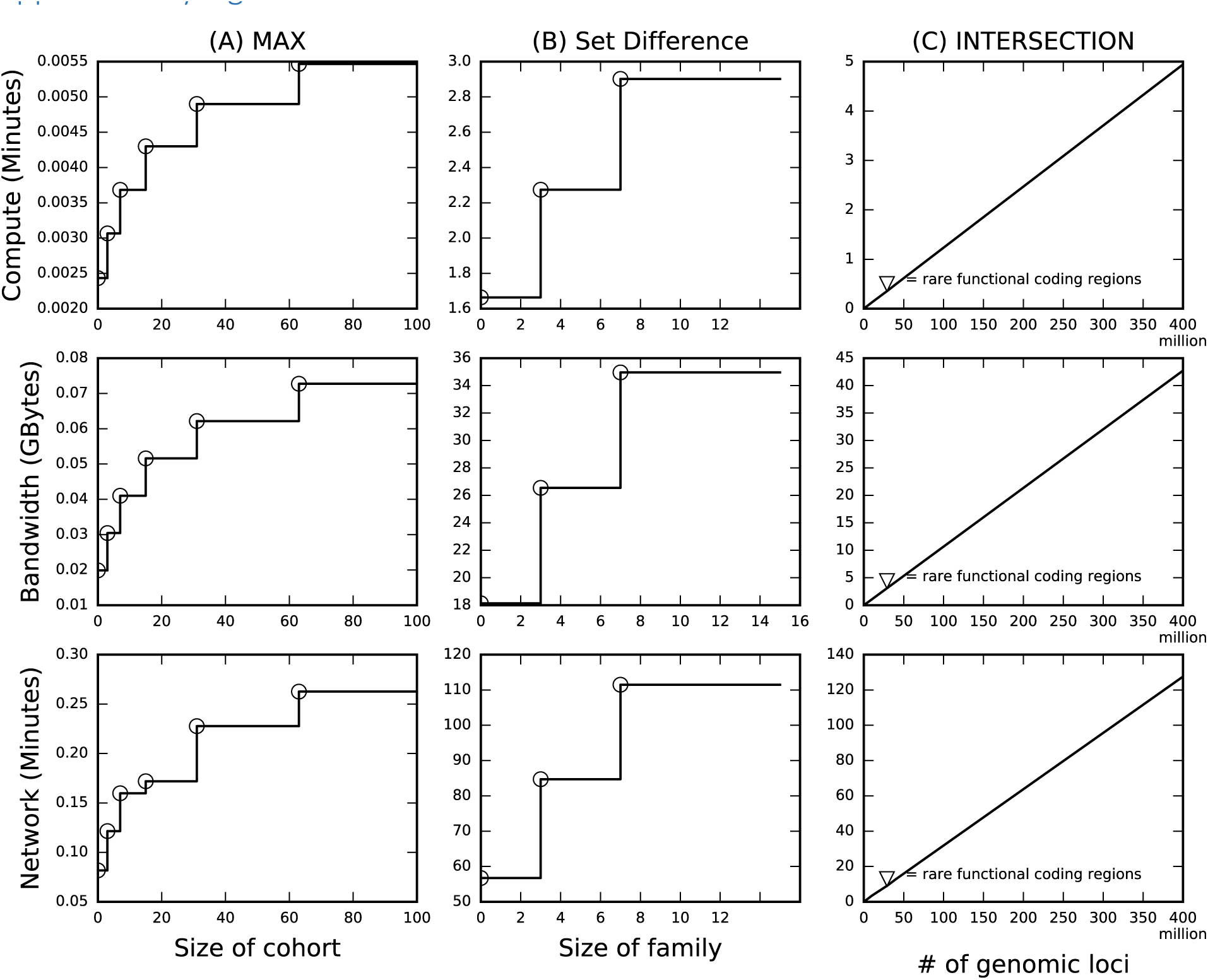
Performance scale up for secure computation. Bandwidth, compute (CPU) time, and overall protocol execution (wall clock) time for the secure MAX, SETDIFF and INTERSECTION scenarios of Table 1, using a single thread on two servers, one located on the East Coast and the other on the West Coast. (A) When increasing the number of unrelated subjects in a small cohort study, all parameters grow logarithmically. (B) When increasing the number of family members in an affected / non affected scenario, parameters also grow logarithmically. (C) In the two hospital scenario, when increasing the number of genomic positions of potential interest (e.g., from the exome to the non-coding genome), all parameters grow linearly. Note that all three scenarios (A-C) perform the bulk of their computation on each element of the input vector separately (Supplementary Figure 1c). All scenarios are thus simple to parallelize for maximum speed-up using multiple threads and nodes.

**Supplementary Figure 5.**
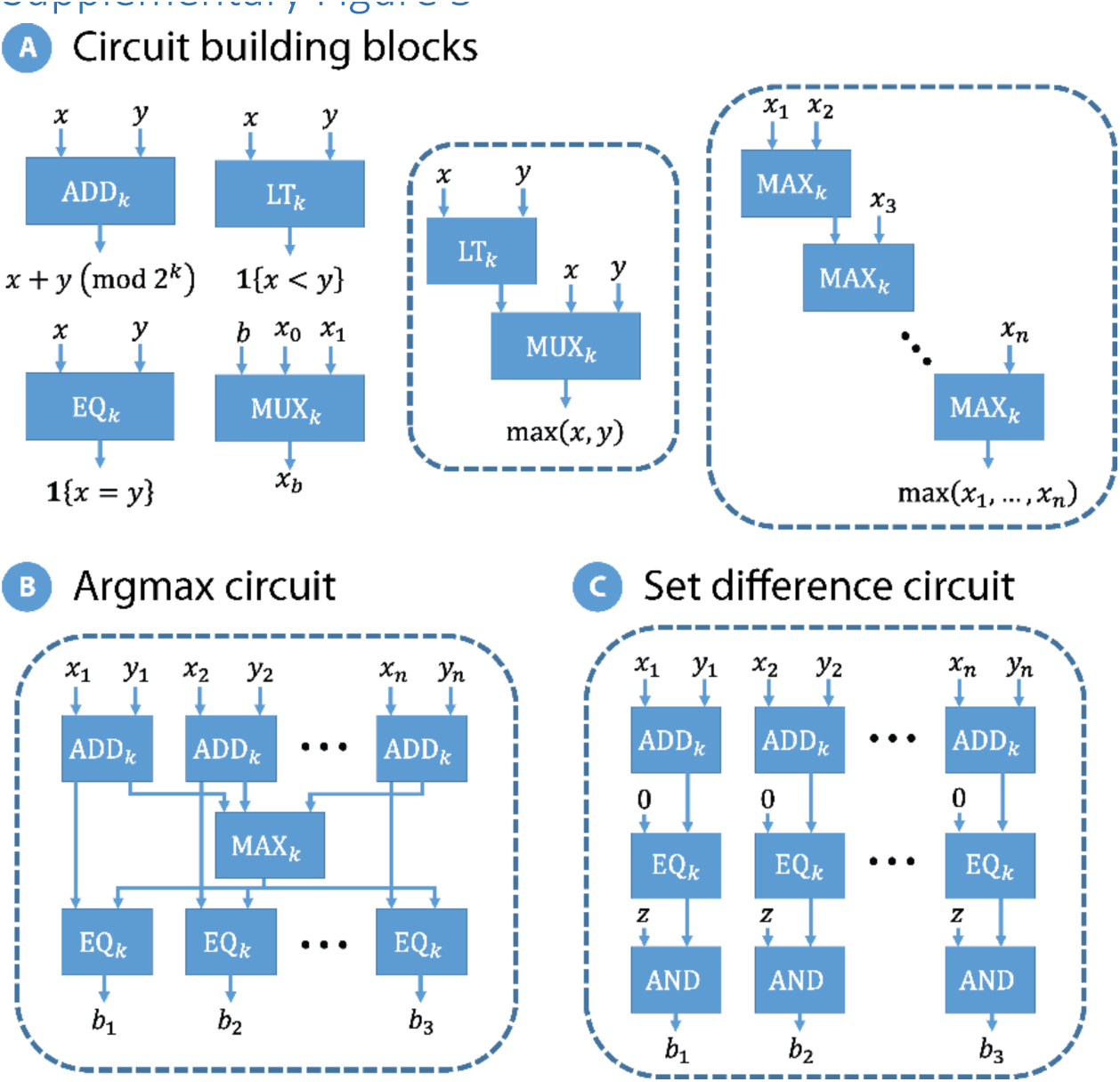
Boolean building blocks for composing complex functions. (A) The basic building blocks we use to build our circuits for identifying common mutations and shared de novo variants include addition, comparison, equality, and multiplexer circuits. An addition circuit ADD_*k*_ on *k* bit inputs takes two *k*-bit values and outputs the *k*-bit representation of their sum (addition is performed modulo 2^*k*^). The LT_*k*_ and EQ_*k*_ circuits implement the less-than and equality operations, respectively, on *k*-bit inputs. The MUX_*k*_ circuit implements a multiplexer circuit which on inputs a selection bit *b* ∈ {0,1} and two *k*-bit values *x*_0_, *x*_1_, outputs *x_b_*. The individual circuits can be efficiently constructed using AND gates and XOR gates, as described by Kolesnikov et al^41^. These basic circuit building blocks can be composed to build a max circuit MAX_*k*_ on two inputs (each of length *k*), which in turn can be used to build a max circuit on *n* inputs. (B) This circuit computes the argmax over *n* additively secret-shared values *z*_1_, …, *z_n_*. The circuit operates by first combining the shares and then taking the max over the resulting vector of values. The argmax is represented by a bit-string of length *n*, where a position *i* has value 1 if *z_i_* is equal to the max value, and 0 otherwise. (C) This circuit computes the set of genes (represented by indices) that are present in a test vector, *z*_1_, …, *z_n_* but not present in a pool *t*_1_, …, *t_n_*. The counts of the mutations appearing in the pool are additively secret shared. The circuit first combines the shares, and then identifies the indices *i* that appear in the test vector (*z_i_* = 1), but not present in the pool (*t_i_* = 0). The circuit outputs *b_i_* = 1 if the gene indexed by *i* occurs in the target genome but not in the test pool, and 0 otherwise.

## References

1 Yang, Y. et al. Clinical Whole-Exome Sequencing for the Diagnosis of Mendelian Disorders. N. Engl. J. Med. 369, 1502–1511 (2013).

2 Iglesias, A. et al. The usefulness of whole-exome sequencing in routine clinical practice. Genet. Med. 16, 922–931 (2014).

3 Lee H, Deignan JL, Dorrani N & et al. CLinical exome sequencing for genetic identification of rare mendelian disorders. JAMA 312, 1880–1887 (2014).

4 Rehm, H. L. et al. ACMG clinical laboratory standards for next-generation sequencing. Genet. Med. Off. J. Am. Coll. Med. Genet. 15, 733–747 (2013).

5 Lek, M. et al. Analysis of protein-coding genetic variation in 60,706 humans. Nature 536, 285–291 (2016).

6 Ng, S. B. et al. Targeted capture and massively parallel sequencing of 12 human exomes. Nature 461, 272–276 (2009).

7 Ng, S. B. et al. Exome sequencing identifies the cause of a mendelian disorder. Nat. Genet. 42, 30–35 (2010).

8 Ng, S.B. et al. Exome sequencing identifies MLL2 mutations as a cause of Kabuki syndrome. Nat. Genet. 42, 790–793 (2010).

9 Moreno-Estrada, A. et al. The genetics of Mexico recapitulates Native American substructure and affects biomedical traits. Science 344, 1280–1285 (2014).

10 Mailman, M.D. et al. The NCBI dbGaP database of genotypes and phenotypes. Nat. Genet. 39, 1181–1186 (2007).

11 Siu, L.L. et al. Facilitating a culture of responsible and effective sharing of cancer genome data. Nat. Med. 22, 464–471 (2016).

12 Shringarpure, S.S. & Bustamante, C. D. Privacy Risks from Genomic Data-Sharing Beacons. Am. J. Hum. Genet. 97, 631–646 (2015).

13. Regalado, A. Networks of Genome Data Will Transform Medicine. MIT Technology Review Available at: https://www.technologyreview.com/s/535016/internet-of-dna/. (Accessed: 30th September 2016)

14 Wagner, J., Paulson, J. N., Wang, X., Bhattacharjee, B. & Corrada Bravo, H. Privacy-preserving microbiome analysis using secure computation. Bioinformatics 32, 1873–1879 (2016).

15 Simmons, S., Sahinalp, C. & Berger, B. Enabling Privacy-Preserving GWASs in Heterogeneous Human Populations. Cell Syst. 3, 54–61 (2016).

16 Popic, V. & Batzoglou, S. Privacy-Preserving Read Mapping Using Locality Sensitive Hashing and Secure Kmer Voting. bioRxiv 046920 (2016). doi: 10.1101/046920

17 Lindell, Y. & Pinkas, B. Secure Multiparty Computation for Privacy-Preserving Data Mining. J. Priv. Confidentiality 1, 59–98 (2009).

18 Yao, A.C.-C. Protocols for Secure Computations. in Annual Symposium on Foundations of Computer Science 160–164 (1982).

19 Simpson, M. A. et al. Mutations in NOTCH2 cause Hajdu-Cheney syndrome, a disorder of severe and progressive bone loss. Nat. Genet. 43, 303–305 (2011).

20 Rivière, J.-B. et al. De novo mutations in the actin genes ACTB and ACTG1 cause Baraitser-Winter syndrome. Nat. Genet. 44, 440–444, S1–2 (2012).

21 McBride, K.L. et al. NOTCH1 mutations in individuals with left ventricular outflow tract malformations reduce ligand-induced signaling. Hum. Mol. Genet. 17, 2886–2893 (2008).

22 Fenske, S. et al. HCN3 contributes to the ventricular action potential waveform in the murine heart. Circ. Res. 109, 1015–1023 (2011).

23 Lindell, Y. & Pinkas, B. A Proof of Security of Yao’s Protocol for Two-Party Computation. J Cryptol. 22, 161–188 (2009).

24 Hazay, C. & Lindell, Y. Efficient Secure Two-Party Protocols - Techniques and Constructions. (Springer, 2010).

25 Dwork, C. Differential Privacy. in ICALP 1–12 (2006).

26 Dinur, I. & Nissim, K. Revealing information while preserving privacy. in PODS 202–210 (2003).

27 Li, H. & Durbin, R. Fast and accurate long-read alignment with Burrows-Wheeler transform. Bioinforma. Oxf. Engl. 26, 589–595 (2010).

28 McKenna, A. et al. The Genome Analysis Toolkit: a MapReduce framework for analyzing next-generation DNA sequencing data. Genome Res. 20, 1297–1303 (2010).

29 Wang, K., Li, M. & Hakonarson, H. ANNOVAR: functional annotation of genetic variants from high-throughput sequencing data. Nucleic Acids Res. 38, e164–e164 (2010).

30 .Cunningham, F. et al. Ensembl 2015. Nucleic Acids Res. 43, D662–669 (2015).

31 Goldreich, O., Micali, S. & Wigderson, A. How to Play any Mental Game or A Completeness Theorem for Protocols with Honest Majority. in Annual ACM Symposium on Theory of Computing 218–229 (1987).

32 Katz, J. & Lindell, Y. Introduction to Modern Cryptography. (Chapman and Hall/CRC Press, 2007).

33 Rabin, M. O. How To Exchange Secrets with Oblivious Transfer. IACR Cryptol. EPrint Arch. 2005, 187 (2005).

34 Kilian, J. Founding Cryptography on Oblivious Transfer. in Proceedings of the 20th Annual ACM Symposium on Theory of Computing, May 2-4, 1988, Chicago, Illinois, USA 20–31 (1988).

35 Naor, M. & Pinkas, B. Oblivious Transfer and Polynomial Evaluation. in Proceedings of the Thirty-First Annual ACM Symposium on Theory of Computing, May 1-4, 1999, Atlanta, Georgia, USA 245–254 (1999).

36 Bellare, M., Hoang, V. T., Keelveedhi, S. & Rogaway, P. Efficient Garbling from a Fixed-Key Blockcipher. in IEEE Symposium on Security and Privacy 478–492 (2013).

37 Kolesnikov, V. & Schneider, T. Improved Garbled Circuit: Free XOR Gates and Applications. in International Colloquium on Automata, Languages and Programming 486–498 (2008).

38 Zahur, S., Rosulek, M. & Evans, D. Two Halves Make a Whole - Reducing Data Transfer in Garbled Circuits Using Half Gates. in EUROCRYPT 220–250 (2015).

39 Damgard, I., Pastro, V., Smart, N. P. & Zakarias, S. Multiparty Computation from Somewhat Homomorphic Encryption. in CRYPTO 643–662 (2012).

40 Ben-Or, M., Goldwasser, S. & Wigderson, A. Completeness Theorems for Non-Cryptographic Fault-Tolerant Distributed Computation (Extended Abstract). in STOC 1–10 (1988).

41 Kolesnikov, V., Sadeghi, A.-R. & Schneider, T. Improved Garbled Circuit Building Blocks and Applications to Auctions and Computing Minima. in Cryptology and Network Security 1–20 (2009).

42 Asharov, G., Lindell, Y., Schneider, T. & Zohner, M. More efficient oblivious transfer and extensions for faster secure computation. in ACM CCS 535–548 (2013).

